# Improving Generalizability in Whole-Cell Antibiotic Discovery Through Active Learning

**DOI:** 10.64898/2026.07.04.736489

**Authors:** Lia R. Serrano, Andrew Zhou, Ziming Wei, Kee-Lee K Stocks, Yasha Ektefaie, Peter J Gwynne, Eric Chen, Inna Krieger, James Sacchettini, Bree Aldridge, Linden T Hu, Maha Farhat

## Abstract

Machine learning (ML) has accelerated molecular discovery, yet training models to generalize to out-of-distribution (OOD) chemical spaces remains fundamentally constrained by the high cost of experimental validation. In antibiotic discovery, where whole-cell phenotypic high throughput screening (HTS) is resource-intensive, iterative ML-guided compound selection – or Active Learning (AL) – offers a pathway to efficiently navigate available chemical spaces. However, the algorithmic tradeoffs between prioritizing compound novelty (exploration), predicted bioactivity (exploitation), and their impact on OOD generalizability remain unresolved for noisy, whole-cell biological systems. In this work, we systematically evaluate three AL strategies for whole-cell bacterial bioactivity and benchmark their effects on model accuracy, hit rate, and OOD performance. Using retrospective simulations on *Mycobacterium tuberculosis* HTS data, we identify an optimal AL strategy that balances predicted hit/non-hit novelty with overall hit rate. We then integrate the strategy in a closed-loop *Borrelia burgdorferi* antibiotic discovery HTS campaign. The AL-guided approach successfully increased the experimental screening hit rate five-fold (from a 0.2% rate within investigator-selected plates to 1.0%). Further, when the trained model was applied in prospective *in silico* selection of highly diverse compounds across multiple bacterial species, the AL-trained whole-cell inhibition predictor demonstrates 53-fold enrichment over investigator-directed screening (11.0% experimental validation of predicted hits). Of these, 100% demonstrated the intended narrow spectrum activity for *Borrelia burgdorferi*. These results demonstrate that calibrated AL strategies can overcome data acquisition bottlenecks and train generalizable property predictors able to extrapolate to OOD molecules.

## Main

Machine learning (ML) promises to reshape molecular discovery and enable the rapid *in silico* screening of the estimated chemical space of 10^60^ drug-like small molecules.^1^ However, the success of molecular property ML models remains constrained by their reliance on static, pre-existing training datasets often acquired for other purposes than model training. Consequently, these models frequently fail to generalize to out-of-distribution (OOD) regions of chemical space, yielding high false-positive and -negative rates when exploring novel molecular scaffolds.^2–10^ This algorithmic limitation is especially acute in whole-cell phenotypic screening, such as for antibiotic discovery, where experimental high throughput screening (HTS) is costly and resource intensive. Unlike targeted *in vitro* assays, whole-cell bacterial inhibition models must learn complex, poorly understood structure-activity relationships, such as cell wall permeability, intracellular stability and compensatory cellular efflux mechanisms. This complexity results in extreme experimental class imbalances (discovery hit rates are characteristically low and as low as <0.1%).^11–13^

A promising strategy to overcome the OOD generalization barrier is to tightly integrate ML prediction with prospective experimental screening. Active Learning (AL) is a closed-loop framework wherein samples are iteratively and selectively labeled for supervised model training based on their informational value to the model. While AL has proven effective in steering compound selection for targeted bioactivity tasks, often using *in-silico* oracle-based validation techniques,^13–36^ its application to complex, whole-cell activity prediction for antibiotic development remains largely uncharted. Further, across both targeted and whole-cell activity prediction, an algorithmic challenge in applying AL is the design of the acquisition strategy. The theoretical trade-offs between prioritizing compound novelty (exploration to improve OOD performance) and predicted inhibition (exploitation to discover and learn structure-activity relationships in bioactives) are not well understood and can vary greatly between applications. In fact, uncalibrated sample selection can inadvertently hurt the performance of ML inhibition models^13,37,38^ by prioritizing redundant, uninformative or adversarial OOD compounds. Conversely, an optimal AL strategy promises to drastically reduce upfront experimental costs by charting under sampled domains with maximal efficiency.^14,15,39^

Here, we present an AL strategy designed to overcome data acquisition bottlenecks and optimize OOD generalizability for whole-cell phenotypic screening. We formulate an AL acquisition function, compatible with batched HTS testing that balances molecular novelty, inhibition prediction and intra-plate chemical diversity. We first benchmark the trade-offs of this strategy against two alternative AL strategies in a retrospective simulation of an *M. tuberculosis* antibiotic discovery campaign through HTS (n = 52,040 of 114,933 possible compounds). We then prospectively validate the selected strategy in a real-time, closed-loop HTS antibiotic discovery campaign against *B. burgdorferi,* the causative agent of Lyme disease, and *E. coli* to model commensal bacteria for narrow-spectrum antibiotics discovery. Iteratively screening batches pulled from a library of ∼292K compounds, we demonstrate that our AL strategy increases experimental hit rates by five-fold while nearly doubling model performance (after hit-rate control). Further, we demonstrate the generalizability to a larger chemical space of the AL trained models through a prospective *in silico* screening campaign of 1.2 million compounds, followed by experimental screening of the 73 top candidates by novelty, selectivity, bioavailability and toxicity. The AL trained inhibition predictors demonstrate robust OOD performance (precision difference <0.03% of training hold out vs prospective screening) and results in a 53-fold higher hit rate, and 1.6-fold higher selectivity than expected through random HTS. Ultimately, we present an AL strategy that balances novelty and inhibition to successfully navigate large chemical spaces and discover novel, potent, and selective antibacterials.

## Results

### Benchmarking Acquisition Strategies for OOD Generalization

To systematically assess acquisition strategies in AL for whole-cell biological data, we simulated a retrospective iterative screening campaign using an existing HTS dataset of 114,933 compounds evaluated for *M. tuberculosis* (*Mtb*) growth inhibition (**Figure 1**). We trained an evidential^40^ directed message passing neural network (DMPNN) classifier to predict growth inhibition.^41^ To simulate the constraints of vendor-manufactured plates of small molecules common in HTS, we predefined batch size at 384 compounds and restricted our labeling budget to ∼50% of the total dataset (n = 55,040) drawn from 103,440 compounds eligible for acquisition; 11,493 compounds were reserved as a holdout test set, designated using a k-means split strategy (*Simulated M. tuberculosis Active Learning,* **Methods**).

**Figure 1.**
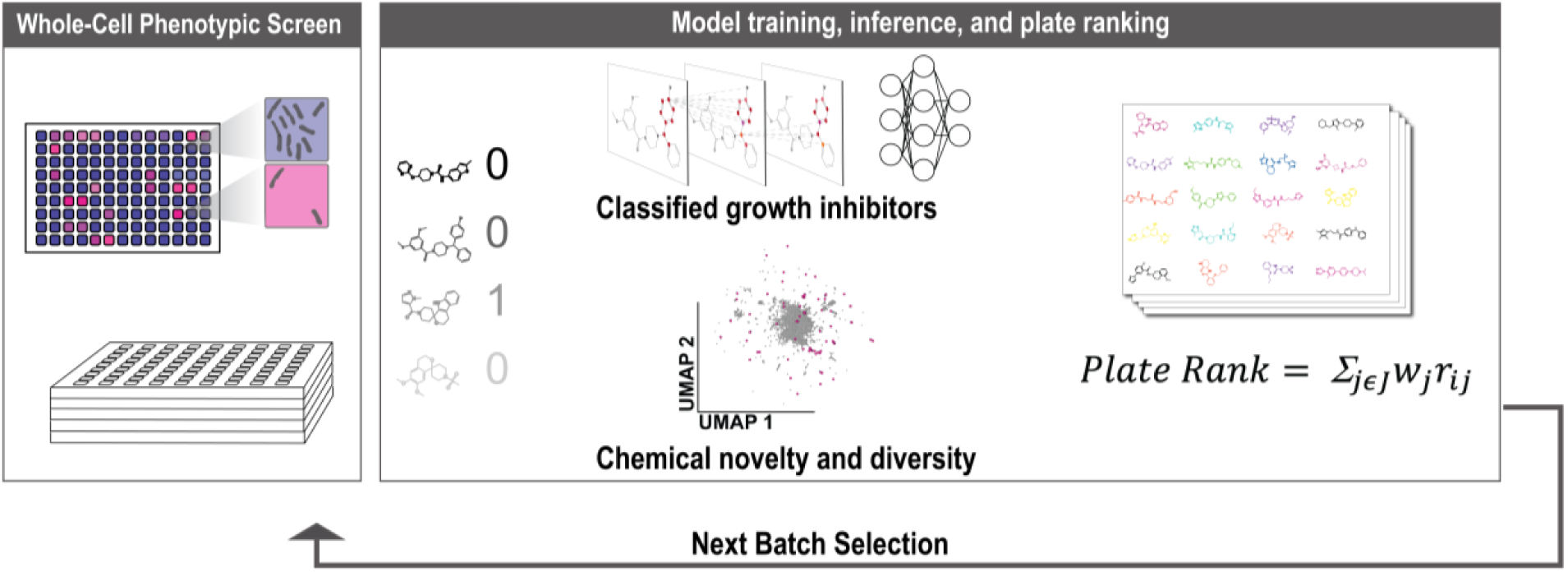
Workflow for prioritizing chemical screening plates in a simulated Active Learning benchmark applied to structure-activity modeling for M. tuberculosis whole-cell inhibition. Binarized growth inhibition data are used to iteratively train an evidential directed message passing neural network. Compounds assigned to virtual plates are ranked based on one to several metrics (*Simulated M. Tuberculosis Active Learning,* Methods). Rankings are computed into an aggregate plate-based ranking which determines the next batch within the AL loop.

We first evaluated two distinct AL acquisition functions against a random selection baseline: an “Inhibition-only” strategy (pure exploitation, prioritizing molecules with the highest probability of growth inhibition), and a “Diversity-Novelty-Inhibition” strategy (balancing exploitation with exploration by prioritizing intra-batch diversity and chemical uniqueness from past iterations, *Simulated M. Tuberculosis Active Learning,* **Methods**). Our simulations demonstrate that both AL schemes significantly boost hit discovery relative to random selection (by up to 11% per iteration, **Figure 2A, 2B**). However, the Diversity-Novelty-Inhibition scheme outperforms Inhibition-only by maintaining a consistently more favorable precision-recall trade-off for models trained on more than ∼ 40,000 compounds (**Figure 2C**).

**Figure 2.**
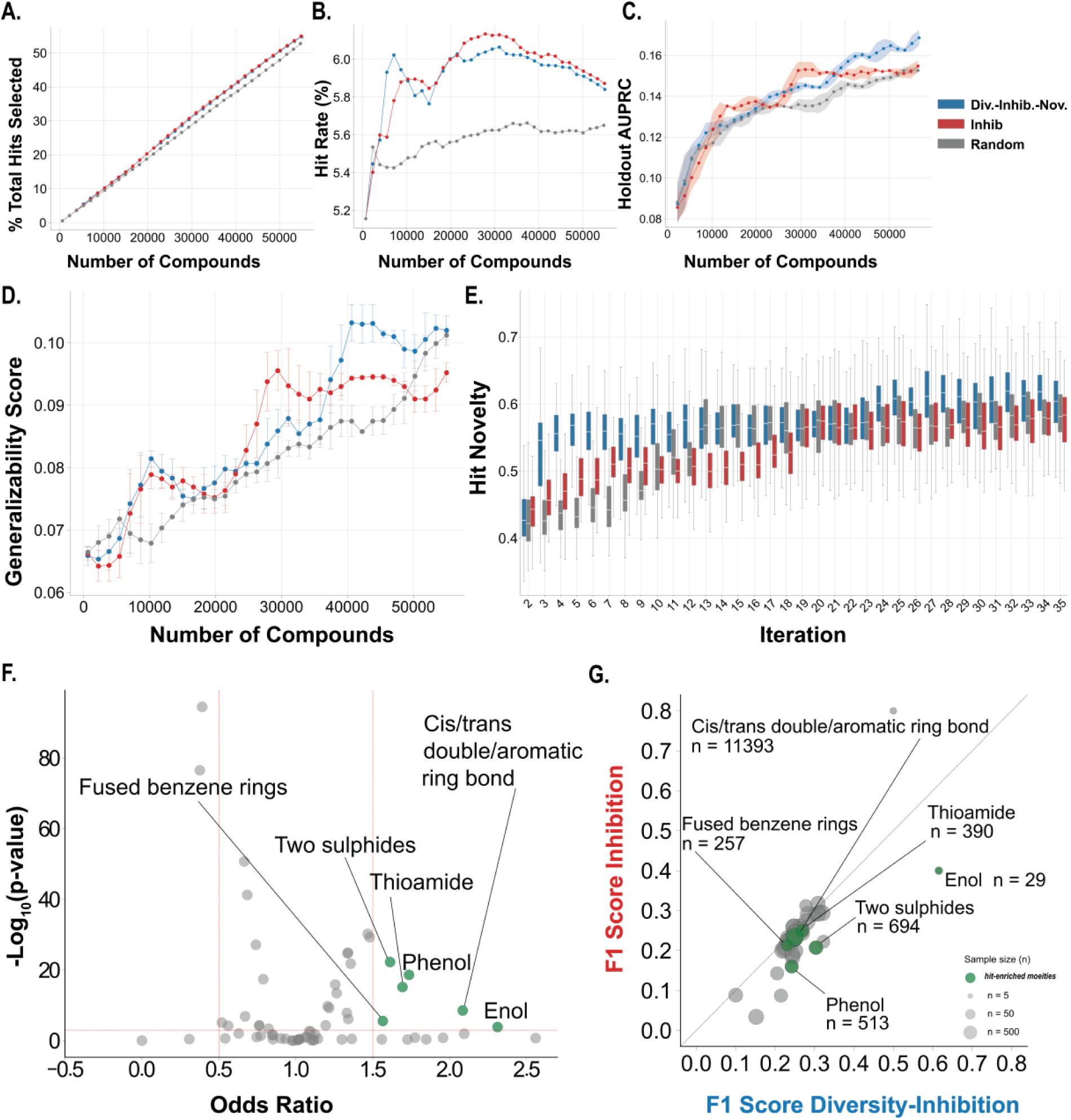
Benchmarking of three AL strategies on *Mtb* HTS data. **(A)** Percent total hits screened vs number of compounds **(B)** Number of active compounds screened per number of molecules screened (“hit rate”). **(C)** AUPRC vs number compounds. Each datapoint represents a new training batch selected based on the Random (gray), Diversity-Inhibition-Novelty (blue), or Inhibition (red) acquisition functions. **(D)** Generalizability score (Methods) vs. number compounds for each condition. **(E)** Novelty (one minus the min. cosine similarity between MiniMol fingerprint of new hit and each hit from previous iteration) of active compounds screened at each iteration relative to the previous iteration. **(F)** Fisher’s exact test was used to test for significant enrichment of medicinally relevant chemical moieties in *M. tuberculosis* inhibitors (n = 6,572). Significantly-enriched chemical moieties (p-value ≤ 0.001, odds ratio ≥ 1.5) are labeled and color-coded in green. **(G)** Performance on holdout molecules summarized by F1 score split by membership in 50 medicinally relevant chemical moieties for the Inhibition-only vs Diversity-Inhibition-Novelty models are plotted. Chemical moieties enriched in hits from our dataset are color-coded in green.

To quantify the impact of AL on OOD generalizability, we computed a Generalizability Score (*Generalizability Score Calculation***, Methods**) for each iteration of our simulation using a holdout set of 11,493 compounds. **Figure 2D** demonstrates that the Diversity-Novelty-Inhibition strategy generalized significantly more accurately than random selection, or pure exploitation. Furthermore, while AL strategies uncover hits at a comparable absolute rate, the Diversity-Novelty-Inhibition strategy demonstrated more efficient identification of novel hits requiring significantly fewer AL iterations to plateau in hit novelty (**Figure 2E**). The Diversity-Novelty-Inhibition strategy also demonstrates superior precision and recall of compounds carrying substructures enriched among *Mtb* inhibitors than the Inhibition-only strategy. This suggests superior learning of relevant structure-activity relationships (**Figure 2F, G**).

### Evaluating the Impact of Evidential Uncertainty in Compound Selection

Incorporating model uncertainty is a theoretically powerful strategy for AL.^22,42–51^ We integrated Dirichlet-based uncertainty (**Eqn. 2**) and aleatoric uncertainty (**Eqn. 3**) into our acquisition function, generating a “Diversity-Novelty-Inhibition-Uncertainty” strategy (*Simulated M. Tuberculosis Active Learning,* **Methods**). This approach explicitly prioritizes compounds for which evidential DMPNN lacks relevant data for inhibition prediction.

Across three metrics –percent of hits selected, overall hit rate, and global AUPRC, Diversity-Novelty-Inhibition-Uncertainty performs similarly to the other AL strategies and is superior to random screening (**Supplementary Figure S1A-C**). Unsurprisingly, because the uncertainty acts as an additional exploration term, the novelty of the hits discovered at early AL iterations was notably higher than that of Diversity-Novelty-Inhibition (**Supplementary Figure S1D**). However, the uncertainty-aware framework performed similarly to random selection on generalizability (**Supplementary Figure S1E**). To further understand this observation, we examined the average minimum cosine similarity of train molecules relative to the remaining selection pool as a proxy for “Remaining Library Novelty”, showing that although the uncertainty-based function effectively selects novel hits relative to train hits, it explores non-hit novelty less effectively than Diversity-Novelty-Inhibition (**Supplementary Figure S1F**). This can be explained by the higher average Dirichlet uncertainty computed for hits compared to non-hits across iterations (percent increase between 1% and 15%). Further, we suspected that the less effective non-hit exploration could be due to the poor calibration of the uncertainty metric in early iterations of AL. Indeed, we see several early AL iterations wherein the median ratio of accurate class evidence to total evidence is below 0.5. This results in an underestimation of uncertainty, rendering this metric unreliable for early AL.

### Prospective Closed-Loop Al Resolves Extreme Class Imbalance in Real-Time HTS

We next transitioned from retrospective simulation to a prospective, real-time closed-loop campaign performed sequentially over ∼1.5 years (**Figure 3A**). We targeted the discovery of antibiotics against *B. burgdorferi* - that spare commensal *E. coli*. Both bacteria differ from *M. tuberculosis* in having a substantially lower random hit rate in HTS (∼0.04% and ∼0.2% for *E. coli* and *B. burgdorferi*, respectively, vs. 1-5% for *M. tuberculosis*) (**Figure 3B**).

**Figure 3.**
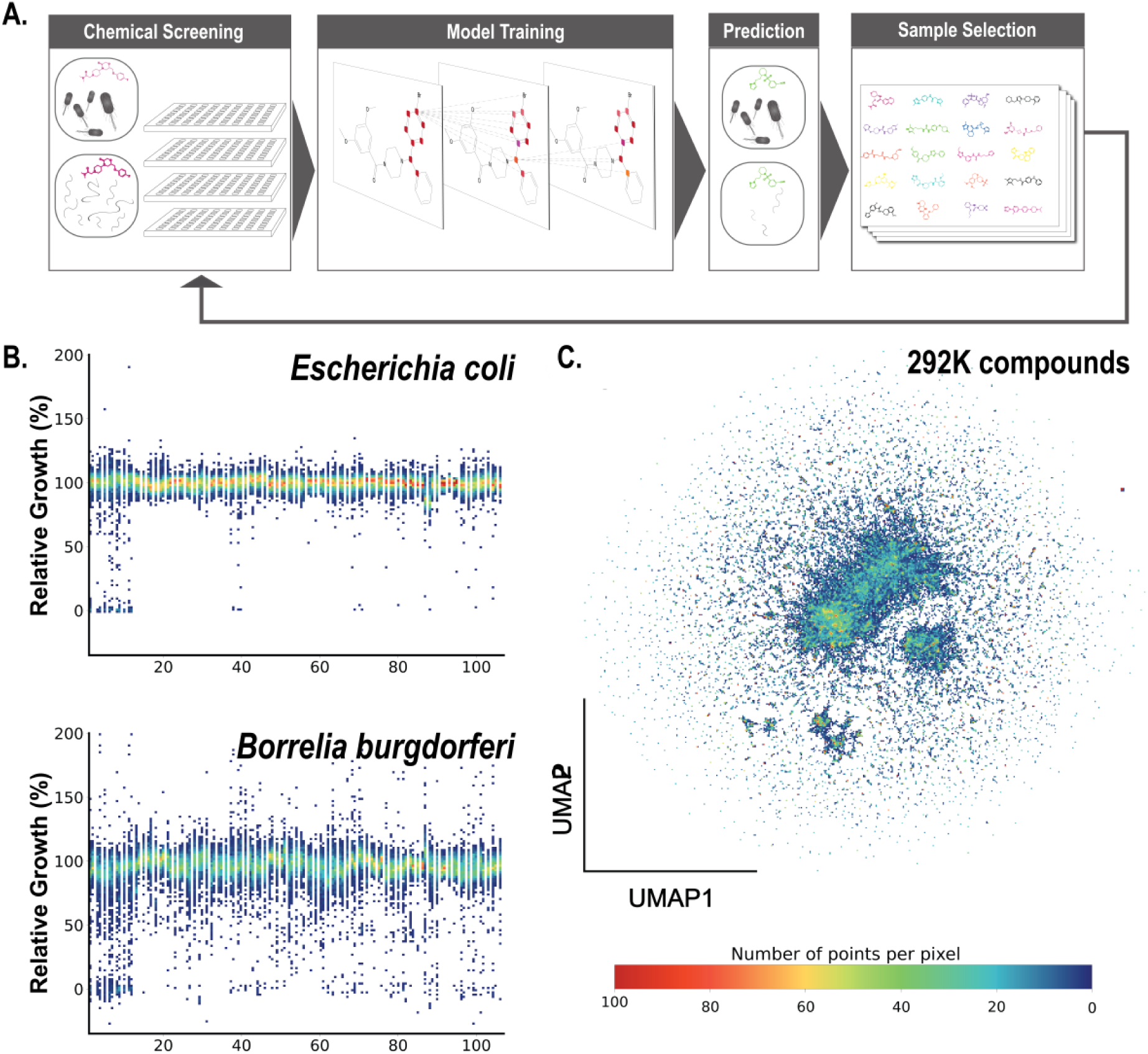
Active Learning Guided Antibiotic Discovery. **(A)** Narrow-spectrum active learning pipeline starting with dual *E. coli* and *B. burgdorferi* high throughput compound screens. Successive screens are selected from a library of 281,459 molecules based upon DMPNN and molecular fingerprint-based model predictions and other data valuation metrics. **(B)** Distribution of active compounds against *E. coli* and *B. burgdorferi* across 384-well plates. **(C)** UMAP projection of 512-dimensional representation (MiniMol) of chemical compound candidates for AL-based HTS.

To initialize AL, we pre-trained a DMPNN with a Dirichlet head on a curated library of 10,422 diverse, bioactive, and/or bioavailable compounds (*Real-time B. burgdorferi Active Learning***, Methods**). We then deployed the Diversity-Novelty-Inhibition acquisition function to prospectively select plates for HTS from a pool of 890 plates containing 281,459 uncharacterized compounds (**Figure 3C**). Across the eleven subsequent training iterations, the top 5% of plates were selected by the AL model and experimentally tested for inhibiting *B. burgdorferi* (batch sizes ranging from 1,760 to 3,500). To rigorously benchmark the AL, batches five through eleven included non-AL guided plates (either randomly selected or investigator-selected) sizes ranging from 1,760 to 6,000 compounds. All experimental screening results were continuously fed back into the model to close the AL loop.

AL increased 384-well plate hit rate by five-fold to 1.0% from 0.2% delivered by the non-AL guided selection (**Figure 4A-C**). In contrast to the non-AL guided plates, the AL-selected plates were enriched in chemical substructures seen in experimentally validated hits (**Figure 4D-F**).

**Figure 4.**
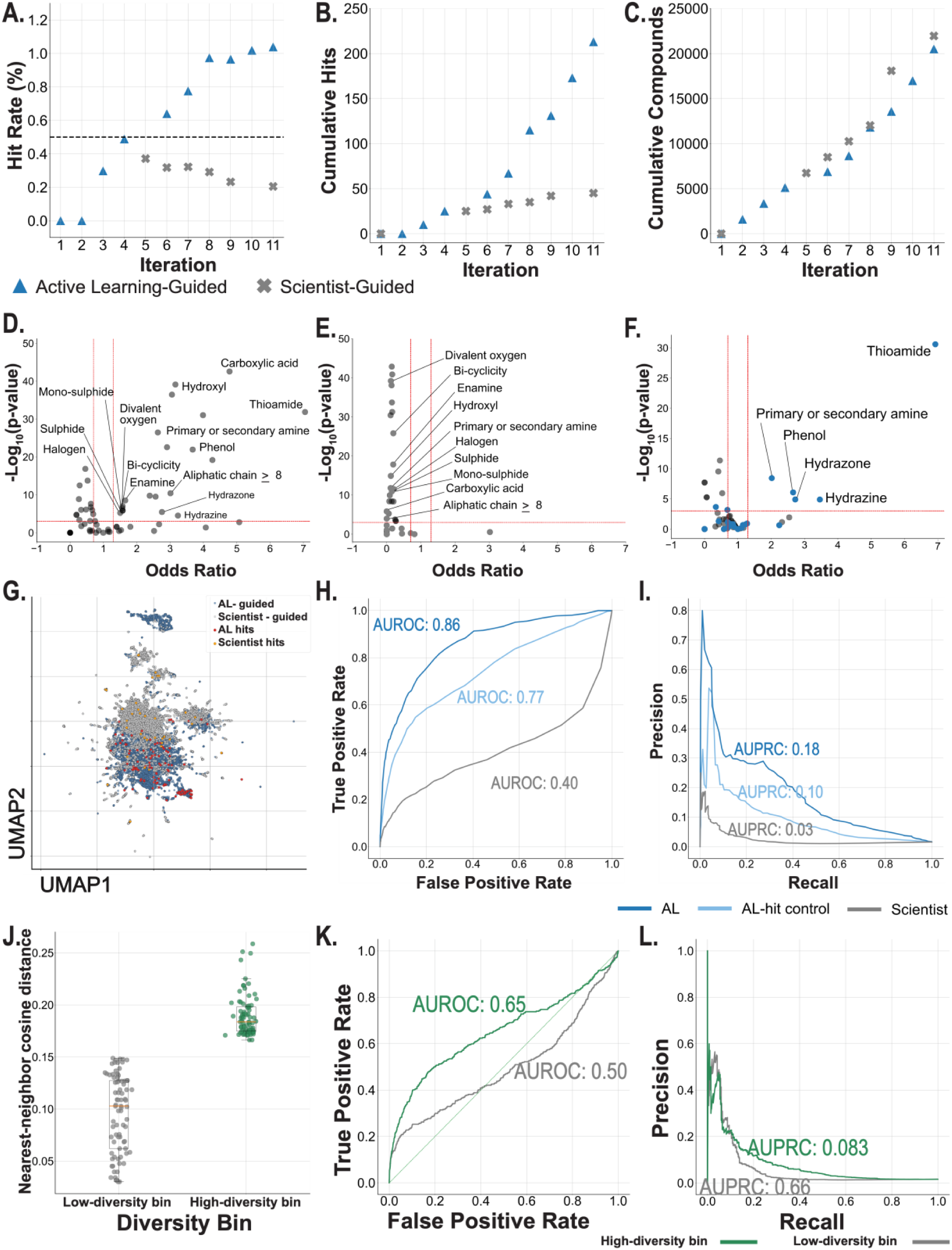
Active Learning-guided sample selection across iterations. Percent hit rate **(A)**, cumulative hits **(B)**, and cumulative compounds **(C)** per model training batch are compared between AL- vs. investigator-guided compound selection. **(D)** Negative log_10_-transformed Fisher’s exact test p-values testing enrichment of medicinally relevant chemical moieties within 578 *B. burgdorferi* growth inhibitors vs. odds ratio. Significantly enriched moieties (p-value ≤ 0.001, odds ratio ≥ 1.5) are labeled. **(E)** Enrichment analysis of medicinally relevant chemical moieties in investigator-selected plates and **(F)** AL-selected plates. Moieties enriched within the hits are colored blue. **(G)** UMAP of MiniMol representations of AL-selected non-hits (blue), investigator-selected non-hits (grey), AL-selected hits (red), investigator-selected hits (orange). **(H)** ROCs for AL, AL-hit control, and investigator-only curated train sets (AUROC 0.86, 0.77, and 0.40, respectively). **(I)** PRCs for AL, AL-hit control, and investigator-only curated train sets (AUPRC 0.18, 0.10, and 0.03, respectively). **(J)** Nearest neighbor cosine distances of hits from, “Low-diversity bin” (grey) and “High-diversity bin” (green) **(K)** ROC of train set with high diversity hits (green) overlaid with ROC of train set with low diversity hits (grey). Each train set has an identical set of non-hits. **(L)** PRC of train set with high diversity hits (green) overlaid with ROC of train set with low diversity hits (grey)

### AL improves Model Performance Independent of Hit-Rate Enrichment

To isolate the algorithmic benefit of AL from AL’s benefits in hit-rate enrichment, we performed an ablation study. We re-trained three distinct models using similar sized datasets: (1) Plates selected purely via the AL scheme (n = 13,469, hit rate = 1.0%), (2) “non-AL” set, i.e. plates selected by human investigators (n = 13,496, hit rate = 0.3%), and (3) And “AL-hit control” set, wherein AL-selected plates were used but true hits were randomly down sampled until the hit rate artificially matched the investigator’s 0.3% baseline (n = 13,376) (**Supplementary Figure S2**). Note, we reserved 25,504 compounds (hits = 404) from our full dataset for a holdout test set.

The model trained on the pure AL data achieved an AUROC of 0.86 ± 0.034 and an AUPRC of 0.18 ± 0.022. In contrast, the non-AL dataset yields an AUROC of only 0.40 ± 0.026 and an AUPRC of 0.03 ± 0.0019. Most notably, when AL-selected data was artificially restricted to the 0.3% non-AL hit rate (AL-hit control), model performance was still substantially higher than the non-AL-trained model (average AUROC across three models trained with different subsampled hit sets = 0.77 ± 0.04, AUPRC = 0.10 ± 0.04) (**Figure 4G-I**). Each of the models tested in this study represent only 25% of our ultimate train set and 4.5% of the total molecules eligible for screening, highlighting the achievable performance gains offered by AL in low data regimes.

To isolate the impact of hit novelty from non-hit novelty and hit rate, we performed an additional study wherein the non-hit set of compounds (n = 26,965, combined AL and non-AL non-hits) was kept constant while we binned a set of 168 hits into two sets of 80 varying in diversity measured by the nearest cosine distance within the set (**Figure 4J**). Using the same holdout set for the previous ablation study, we show that the greater hit diversity resulted in a 25% increase in AUPRC (0.066 ± 0.004 for the lowest diversity and 0.083 ± 0.006 for the highest diversity, respectively) and a 31% increase in AUROC (0.50 ± 0.04 and 0.65 ± 0.02, respectively) (**Figure 4K-L**). These results demonstrate that AL improves model performance through increasing the hit rate as well as through rational selection of both hits and non-hits.

### Virtual Screening and Validation for Narrow-Spectrum Candidates

Using our final models trained on 52,913 compounds (18.1% of the 292K available compounds) (**Supplementary Figure S3**), we virtually screened 1.2M compounds against *B. burgdorferi*. In parallel we built non-AL inhibition prediction models for *S. aureus*, and *E. coli* (**Supplementary Figure S4, Supplementary Table 1**) (**Methods**). *S. aureus* and *E. coli* were selected as representatives of commensal Gram-positive and Gram-negative bacteria, respectively, to support the identification of selective antibiotic candidates for Lyme. We note that 40% of *B. burgdorferi* hits uncovered thus far are nonselective as determined by HTS *E. coli* screening or our *S. aureus* inhibition model possibly because of shared targets (**Figure 5A**).

**Figure 5.**
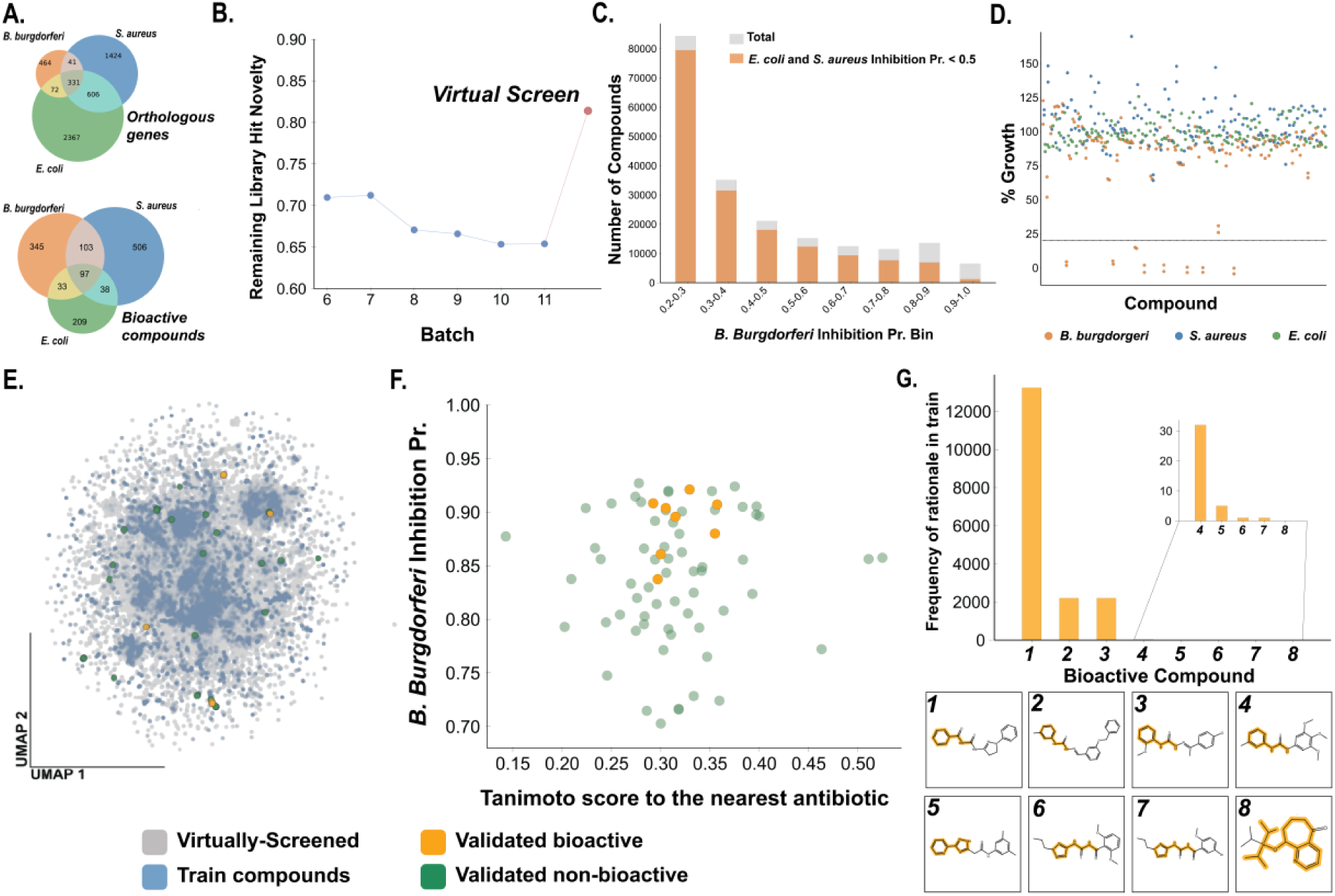
Virtual screen filtering and validation. **A.** Overlap of orthologous genes (top) and bioactive compounds (bottom) across *B. burgdorferi*, *S. aureus*, and *E. coli.* **B.** Proxy metric for Remaining Library Hit Novelty (average minimum cosine similarity of train hits molecules relative to predicted hits from remaining selection pool) for Batches 6-11 and that for the external virtual screening set (red). **C.** Number of total (grey) and selective (orange) compounds across the probability distribution binned in increments of 0.1. **D.** 73 validated compounds percent growth relative to DMSO-treated negative controls. **E.** UMAP of MiniMol representation of train compounds (blue), virtually screened compounds (grey), validated compounds (green), and bioactive compounds from validation experiment (gold). **F.** Model probability of *B. burgdorferi* growth inhibition vs. Tanimoto Similarity score of Morgan Fingerprint representation of each validated compound relative to the nearest antimicrobial entity scraped from chEMBl for non-hits (green) and hits (gold). **G.** Frequency of MCTS-inferred rationales from validated bioactive compounds in train data (indexed 1-8).

We aimed to virtually screen compound libraries to identify antibiotic candidates that were highly novel relative to both our final train set as well as existing antibiotics. The screened compound set (n = 1.2M compounds) has measurable novelty (relative to our train data) compared to the HTS screening pool on which we conducted the AL campaign (**Figure 5B**). Out of our 1.2M compounds, 31,400 were predicted to inhibit *B. burgdorferi* (pr. ≥ 0.7), 15,847 of which were predicted to have < 0.5 probability of inhibiting *E. coli* or *S. aureus* (**Figure 5C**). Of the 15,847 potentially selective *B. burgdorferi* growth inhibitors, we experimentally validated 73 compounds that were within the top 300 molecules ranked based on novelty to existing antibiotics, novelty to the training data, bioavailability, toxicity, and the inhibition probabilities of each organism (*Virtual Screening*, **Methods**).

Eight of the 73 molecules (11%) experimentally inhibited growth by >80%, and 73 of 73 molecules (100%) did not experimentally inhibit *S. aureus* or *E. coli* growth (median inhibition of 0.81% and -24.94% (growth enhancing), respectively **(Figure 5D**). The observed positive predictive value of 11% is consistent with our model’s performance on internal time-split hold out data (PPV = 11%, Recall = 65%, n = 10,540). The experimentally validated hits were highly novel compared with the training set as well as existing antibiotics/antimicrobial compounds scraped from ChEMBL (n = 11,272) with median Tanimoto Similarities of 0.34 and 0.30 to the nearest neighbor in each respective set (**Figure 5E, F**), demonstrating OOD generalizability. To simulate the logical extreme of AL-enabled training efficiency, we tested our model trained only on AL-selected molecules (25.5% of our final training data and 4.5% of the entire screening library) and found that 6 of the 8 true positives were classified as inhibitors, demonstrating 75% recall (**Supplemental Results**). Notably, in addition to the 8 validated hits, 8 additional compounds experimentally inhibited *B. burgdorferi* at levels (at 20-80%, of the control, **Supplementary Figure S5**). We observed a 3.4-fold enrichment in intermediate inhibitors relative to compounds that minimally inhibited growth (<20% relative to control) compared with the training data (Fisher exact P-value = 0.002).

We measured dose dependent inhibition of *B. burgdorferi* of the 8 strong inhibitors identified in the HTS by measuring growth in media across 15 concentrations of each compound (*B. burgdorferi dose response curves***, Methods**). Seven of the eight strong inhibitors had a half maximal inhibitory concentration (IC_50_) < 50 μM, ranging from 5.0 μM to 48.0 μM (**Supplementary Figure S6**). These results confirm that AL trained models predict biologically relevant phenotypic inhibition.

We suspect that there is a minimal active substructure that influenced the inhibitory properties of the strong and intermediate inhibitors. We determined which structural moieties are associated with inhibition using MCTS rationale extraction (*MCTS chemical rationale extraction*, **Methods**) and found one minimal rationale for each of the eight strong inhibitors. One pair of inhibitors shared a rationale, with the remaining six each having a unique rationale. We mined the train molecules for these seven unique substructures to further benchmark model generalizability to OOD molecules. We found that except for one rationale that was present ∼25% of the train molecules, the remaining six occurred in fewer than 7.1% of hits in the training data (4.2% of total train molecules), with one being completely absent from the train set (**Figure 5G**). For the intermediate inhibitors, seven of the eight molecules were found to have minimal rationales, five of which were unique among the combined strong and intermediate inhibitors. Of the five unique rationales, at most, 2.8% of hits (0.9% of total) molecules in the train data contained the same substructure with four being completely absent from all train hits. This analysis underscores our model’s ability to generalize beyond memorized structure-activity relationships to identify compounds with unseen bioactive moieties.

## Discussion

Here, we demonstrate the transition from retrospective static ML to prospective “lab-in-the-loop” discovery. Recent systematic benchmarks have highlighted that molecular discovery is inherently an out-of-distribution (OOD) prediction problem wherein standard ML fails to generalize to novel chemical scaffolds.^52–56^ Our study provides an experimentally validated framework to efficiently select the most informative molecules to improve model performance. By benchmarking Active Learning (AL) acquisition functions retrospectively, we demonstrate that a carefully calibrated strategy that balances predicted inhibition, molecular novelty, and diversity results in models trained with the highest performance on a holdout test set. Our prospective evaluation of this AL strategy on a 281K compound library successfully bypassed standard HTS bottlenecks, achieving a five-fold increase in experimental hit rate and more rapid identification of novel hits ultimately yielding a model with superior OOD generalizability. We also demonstrate that only 4.5% of our AL selection pool was necessary for high-performing structure-activity model training achieving an AUROC and AUPRC of 0.86 and 0.18, respectively, on hold out molecules.

Our findings reveal a dual benefit when AL is applied to noisy, whole-cell phenotypic screening. Because complex biological barriers—such as the restrictive gram-negative-like cell walls of *E. coli* and *B. burgdorferi*—drive empirical hit rates to extreme lows (0.1% to 0.5%), whole-cell inhibition learning suffers from severe class imbalance. Our ablation studies demonstrate that AL can be a dynamic data-augmentation engine. By prioritizing active inhibitors for experimental testing, the algorithm not only accelerated hit identification but the resulting enrichment of hits and their diversity fundamentally improved global model discrimination. In addition, we identified that not all exploration metrics are created equal. The evidential uncertainty metric that we tested was less effective at guiding eventual model discrimination than the more direct novelty metrics to which we compared it. This may be due to the evidential uncertainty being vulnerable to spurious connections between structure and label especially early in training, thus potentially inflating the importance of chemical moieties for a given prediction. Globally our findings confirm the intuition that a well calibrated balance between “exploitation” and “exploration” is needed to position the resultant ML models for extrapolation across a much wider range of OOD structures.

One limitation of our study is that our tested acquisition functions were static and did not adapt to changes in availability of compounds in the untested pool. In the future, a batch-by-batch re-formulation of the optimal selection may further increase efficiency and reduce costs of expensive experimental testing albeit at the likely expense of higher complexity. Our study is also limited by the focus of AL on a single molecular property. Although we demonstrate that single property AL combined with ranking on other properties successfully navigates the chemical space to identify selective *B. burgdorferi* inhibitors i.e. that do not inhibit *E. coli* or *S. aureus*, future work can focus on comparing this approach to AL with multiple simultaneous objectives.

## Conclusion

Our work establishes a new benchmark for practical closed-loop Active Learning in whole-cell antibiotic discovery. We propose a set of metrics that resulted in substantial gains in hit rate of HTS-based antibiotic discovery. AL generated models with better OOD performance and that demonstrate stable generalizability in real-world campaign for identification of selective antibiotics to *B. burgdorferi*. Ultimately, by elegantly bridging the computational-experimental divide, our framework demonstrates how tightly coupled “lab-in-the-loop” AI can transform HTS campaigns into dynamic, highly generalizable engines for therapeutic discovery.

## Methods

### High throughput screening of *M. tuberculosis*

*M. tuberculosis* high throughput screening data was collected according to the protocol described elsewhere.^57^ Growth inhibition was binarized using an internally calibrated threshold based on the distribution of inhibition across the assayed molecules. Molecules with normalized inhibition based on the corresponding plate controls at or more than 40% are labeled as inhibitors, while other molecules are labeled as non-inhibitory. For replicates across different batches, molecules with any replicate that exceed 40% inhibition are labeled as inhibitory.

### *B. burgdorferi* and *E. coli* High throughput screening

*B. burgdorferi* 5A18NP1 containing an exogenous plasmid expressing GFP constitutively was used in this study (kind gift from Drs. Kim Lewis and Nadja Leimer). *B. burgdorferi* was grown at 32° with 2% CO_2_ static in BSK II medium, composed of bovine serum albumin (50.00 g/L), CMRL-1066 [US Biologicals (9.80 g/L)], HEPES (6.60 g/L), peptone (5.60 g/L), dextrose (5.60 g/L), sodium bicarbonate (2.44 g/L), yeast extract (2.20 g/L), sodium pyruvate (1.00 g/L), sodium citrate (0.90 g/L), N-acetyl glucosamine (0.50 g/L),1.4% gelatin, and 6.2% rabbit serum.^58^ The pH was adjusted to 7.6 before the media was filter sterilized and stored at −20°C. All *B. burgdorferi* cultures were grown in the presence of kanamycin 200 μg/mL and streptomycin 50 μg/mL to maintain the GFP plasmid. *Escherichia coli* MG1655 was grown in LB at 37° at 200 rpm shaking. *Staphylococcus aureus* NCTC 8532 (ATCC 12600) was grown in tryptic soy broth (TSB) at 37° at 200 rpm shaking.

All high-throughput screening was completed at ICCB-Longwood Screening Facility at Harvard Medical School with their equipment and compound libraries. 384 well plates (Corning 3701/3764) were filled with 30 μl of media using a Thermo Multipdrop combi. Compounds were added by a pin transfer of 300 nL using a custom-built robot featuring a Peak Robotics KX2-Plus-750 robot arm and Epson T6-B602s SCARA robot. Bacterial cells (30 μL) were then added to the plates using the Thermo Multidrop Combi. *B. burgdorferi* was seeded in a final concentration of 1x10^5^ cells/mL while *E. coli* was seeded in approximately 1:1000 back dilution from an overnight culture. *B. burgdorferi* plates were sealed with aluminum sealing film. *B. burgdorferi* was incubated at 37° 0% CO_2_ for 7 days. GFP signal (ex 485/em 535) and media color (OD_570_) change was measured on a Revvity EnVision plate reader. *E. coli* was incubated at 37° 0% CO_2_ overnight and cell density was measured by OD_600_. Inhibitory compounds were determined by percent inhibition compared to positive and vehicle controls.

A custom compound library containing 73 compounds was purchased from MedChemExpress. Stock compounds were at 10 mM in DMSO. Compounds were transferred into an Echo qualified low dead volume plate using an Agilent Bravo liquid handler. Assay plates were prepared by transferring 300 nL (50 μM final concentration) into 384 well plates (Corning 3701/3764) using a Beckman Coulter Echo 655. *B. burgdorferi, E. coli,* and *S. aureus* were plated and grown as previously described.

### *B. burgdorferi* dose response curves

To perform dose response curves, *B. burgdorferi* GFP was grown and plated using a Thermo Multidrop Combi as described above. Compounds were dispensed in flat bottom 384 well plates (Corning 3764) using a digital drug dispenser (D300e digital dispenser; Hewlett-Packard) for a 15-dose series in 2-fold dilutions with a maximum DMSO percentage of 1%. Doxycycline (1 μg/mL) was used as a positive control for normalization to 100% inhibition.

Data was processed using custom Pythons scripts. GFP signal was normalized to untreated controls for 0% inhibition and doxycycline controls for 100% inhibition. Three biological replicates were fitted to a Hill curve and inhibitory concentrations (ICs) were calculated from the Hill curve.

### Real-time *B. burgdorferi* Active Learning

#### Data

The initial training set consisted of 10,422 compounds from Biomol4, ChemBridge2020, Selleck FDA-Approved 2023, and ChemDiv7. The sampling candidates (n = 281,459) eligible for AL-selection were pre-plated into 890 plates (ICCB-Longwood Screening Facility at Harvard Medical School).

#### Model

We trained a Directed Message Passing Neural Network (DMPNN)^59^ model coupled to a feed-forward network and an evidential Dirichlet head^40^ within Chemprop 1.7.0. This model architecture was selected based upon our previous benchmark of DMPNN with evidential Dirichlet loss against several state-of-the-art architectures for *Mtb* inhibition classification tasks.^57^ We utilized additional learned features^32^ alongside the MPNN representations for the final FFN predictions. The DMPNN has three layers, and the hidden dimension is 300. A Dirichlet loss function was used with an evidential regularization of 0.2. The input data are binarized percent growth inhibition data, requiring at least 80% growth inhibition to be considered active. We applied class balancing within ChemProp to account for the low hit rate. After DMPNN training, the embeddings were concatenated to RDkit fingerprints (Batches 1-7) or MiniMol fingerprints (Batches seven and above).^60,61^ The resulting representations were passed through two feed-forward layers using the ReLU activation function. During training, we implemented the Chemprop one-cycle learning rate schedule (initial rate of 0.0001, maximum rate of 0.001) across 200 epochs. The scaffold split strategy was used to partition train, test, and validation sets (80/10/10) split. The final model is an ensemble of the five replicates.

#### Batch selection

384-well plates are ranked according to the following metrics:

1. Novelty score of compounds with inhibition probability < 0.7
2. Number of novel hits (a molecule within a plate is considered novel if it has a Tanimoto similarity of at most 0.5 relative to each molecule in the previous train set)
3. Number of compounds with inhibition probability > 0.7
4. Number of unique “rationales” (*MCTS chemical rationale extraction*, **Methods**)
5. Number of Butina clusters^62^

Each plate in the compound library receives a ranking for each metric. These ranks are aggregated into a summary rank according to **Eqn. 1**

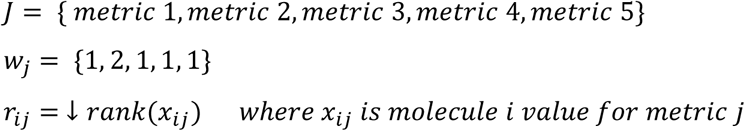

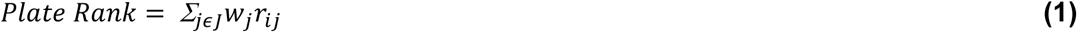

#### MCTS chemical rationale extraction

For a given molecule, we infer the molecular substructure that drives the model’s prediction using a Monte Carlo Tree Search to identify subgraphs of the molecules that are between 8 and 20 atoms with predicted inhibition above 0.7.^63^ This method has been used previously to identify rationales of antibiotic inhibition.^4^ We extract up to 5 rationales per molecule.

#### Generalizability Score Calculation

Generalizability score is computed by partitioning holdout data (n = 114,493) into 10 bins incrementally more chemically divergent from the train set. Chemical divergence is quantified here using the maximum cosine similarity of the MiniMol representations of each test molecule relative to that for the molecules in the training set. The chemical divergence of the hits was computed separately and distributed across bins evenly. The AUPRC of the three most chemically divergent bins is calculated. The area under the AUPRC vs. bin curve is then calculated using the integrate module from Python library SciPy (v. 1.17.1) for each iteration and is termed “Generalizability Score”.

### Simulated *M. tuberculosis* Active Learning

#### Data

The *Mtb* growth inhibition dataset comprised 114,933 distinct molecules, of which 6,572 (5.7%) were annotated as *Mtb* growth inhibitors (compounds or fragments), and 108,361 (94.3%) were annotated as non-inhibitors.

We split the data using the k-means split strategy from Chemprop (v2.2.2) data module to stratify compounds eligible for training (n = 103,440) vs. used as a test set (n = 11,493). Compounds from the *Mtb* dataset were randomly assigned to groups of 320 to emulate the 384-well HTS plate format. For each simulation, the same 2 plates (n = 640 molecules) were used to train the initial model which was used for subsequent AL-based selections.

#### Model

For our retrospective analysis we utilized the same architecture (DMPNN + evidential Dirichlet head) as that used within the real-time Bb AL workflow except there were no additional molecular features appended to the DMPNN embeddings. For each AL train iteration, the model is trained across 75 epochs and the final model is an ensemble of four replicates.

#### Acquisition functions

#### Diversity-Inhibition-Novelty

##### Metrics

1. Novelty score of compounds with inhibition probability < 0.7
2. Number of novel hits (a molecule within a plate is considered novel if it has a Tanimoto Similarity of at most 0.5 relative to each molecule in the previous train set)
3. Number of compounds with inhibition probability > 0.7
4. Number of Butina clusters^62^

Aggregation:

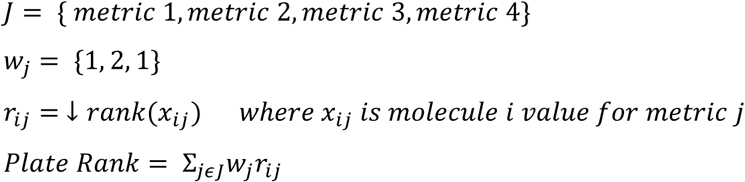

#### Inhibition

Metrics:

1. Average inhibition probability

Aggregation:

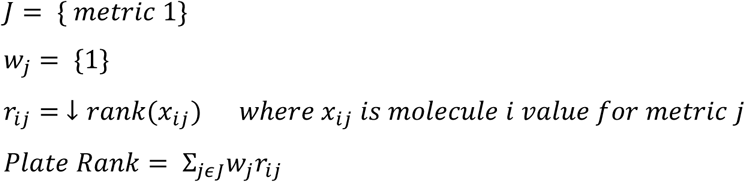

Diversity-Inhibition-Novelty-Uncertainty:

#### Metrics

1. Novelty score of compounds with inhibition probability < 0.7
2. Number of novel hits (a molecule within a plate is considered novel if it has a Tanimoto 2. similarity of at most 0.5 relative to each molecule in the previous train set)
3. Number of compounds with inhibition probability > 0.7
4. Number of Butina clusters
5. Dirichlet Uncertainty 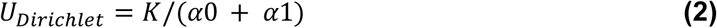
6. Aleatoric uncertainty

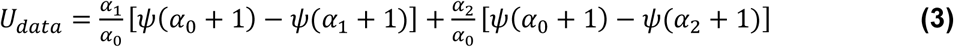

#### Aggregation

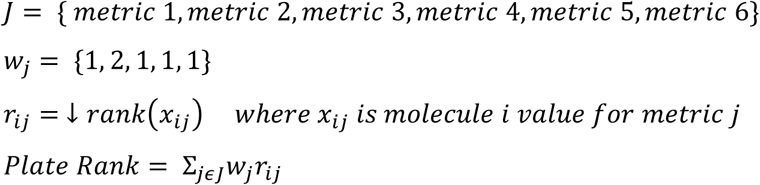

Random:

Virtual plates were randomly selected from the pool of plates not yet used to train the model.

### Virtual Screening

#### E. coli and S. aureus data/model

We trained a DMPNN using the same parameters as described for the *B. burgdorferi* inhibition model (*Real-time B. burgdorferi Active Learning***, Methods**) for both *E. coli* and *S. aureus* inhibition models. For the E. coli model training set, we used the AL batches concurrently tested against *E. coli* and *B. burgdorferi* (n = 52,913) in addition to public datasets, rendering a train set of 154,403 with 475 hits.^2,64–66^ For the *S. aureus* model, we used a public dataset of 327,979 with 1,062 hits.^67^

#### Ranking Strategy

1,203,388 compounds were virtually screened for inhibition against *B. burgdorferi*, *E. coli*, and *S. aureus*. These compounds were filtered to only include compounds with a probability of B. burgdorferi inhibition ≥ 0.7, and probability of *E. coli* and *S. aureus* inhibition ≤ 0.5. Then, we predicted ADMET properties on the subset list.^68^ Each compound was ranked on the following properties:

1. *B. burgdorferi* inhibition (ascending)
2. *E. coli* inhibition (descending)
3. *S. aureus inhibition* (descending)
4. Aggregate Toxicity Score (descending)
5. Novelty to training data (ascending)
6. Novelty to existing antibiotics (ascending)

The Aggregate Toxicity Score was calculated by summing the rank of the individual toxicity metrics (**Supplementary Table 1**) predicted by ADMET-AI and re-ranking the summed value for each compound. Novelty relative to training data was calculated by computing one minus the maximum cosine similarity between the MiniMol-derived representation of a given virtually screened compound and that for the 52,913 training compounds. Novelty relative to existing antibiotics was computed the same way between each virtually screened compound and entities from ChEMBL of known antibiotic activity.^69^ Finally, each of the ranks were summed into an aggregate score which was then re-ranked.

#### Supplemental Results

The AL, AL-hit control, and non-AL models accurately predicted 6, 3, and 1 of the 8 validated hits, respectively. Notably, we see that the AL hit control models are well-calibrated to the chemical space evident in the respective incorrect predictions having higher Dirichlet Uncertainties than the correct predictions. On the other hand, the model trained on investigator-curated data shows low Dirichlet Uncertainty on inaccurate predictions very offset from the binarization threshold (**Supplementary Figure S7**). This implies that the non-AL model over generalizes certain structure-activity relationships due to lack of evidence to the contrary. Curated data used to train the pure AL and AL-hit control model successfully circumvented this pathology common within redundant chemical compound datasets. Further, this result demonstrates the potential sample-labeling efficiency gains of AL applied to whole-cell phenotypic screening. Using only 25.5% of our final dataset and 4.5% of the entire library from which we were selecting samples, we achieve a 75% recall of the bioactive hits from our experimental validation compared to 12.5% recall using the model trained on investigator-curated data. It is important to note that the pure AL train set in the ablation analysis was influenced by the model which was trained on all the available data during the real-time AL selection. However, this experiment affirms that rational train set curation can attain OOD generalizability in low data settings for noisy, plate-based, whole-cell screening assays.

## Acknowledgements

We acknowledge the support of the ICCB-Longwood Screening Facility at Harvard Medical School.

## Author Contributions

LRS, AZ, ZW, KKS, YE methodology; LRS, AZ, ZW, YE software; LRS, AZ, ZW, KKS, IK investigation; LRS, AZ, ZW, KKS, IK data curation; LRS, AZ, ZW, KKS formal Analysis; LRS writing - original draft; LRS, AZ, KKS, ZW, YE, PG, EC, IK, JS, LTH, MRF, BA writing - review & editing; LRS, KKS visualization; LRS, AZ, KKS validation; MRF, LTH, JS resources; MRF, LTH, BA funding acquisition; MRF, LTH, BA supervision; MRF, YE, LTH, BA conceptualization

