## SupplementaryFigures for "Improving Generalizability in Whole-Cell Antibiotic Discovery Through Active Learning"

### Supplementary Figures

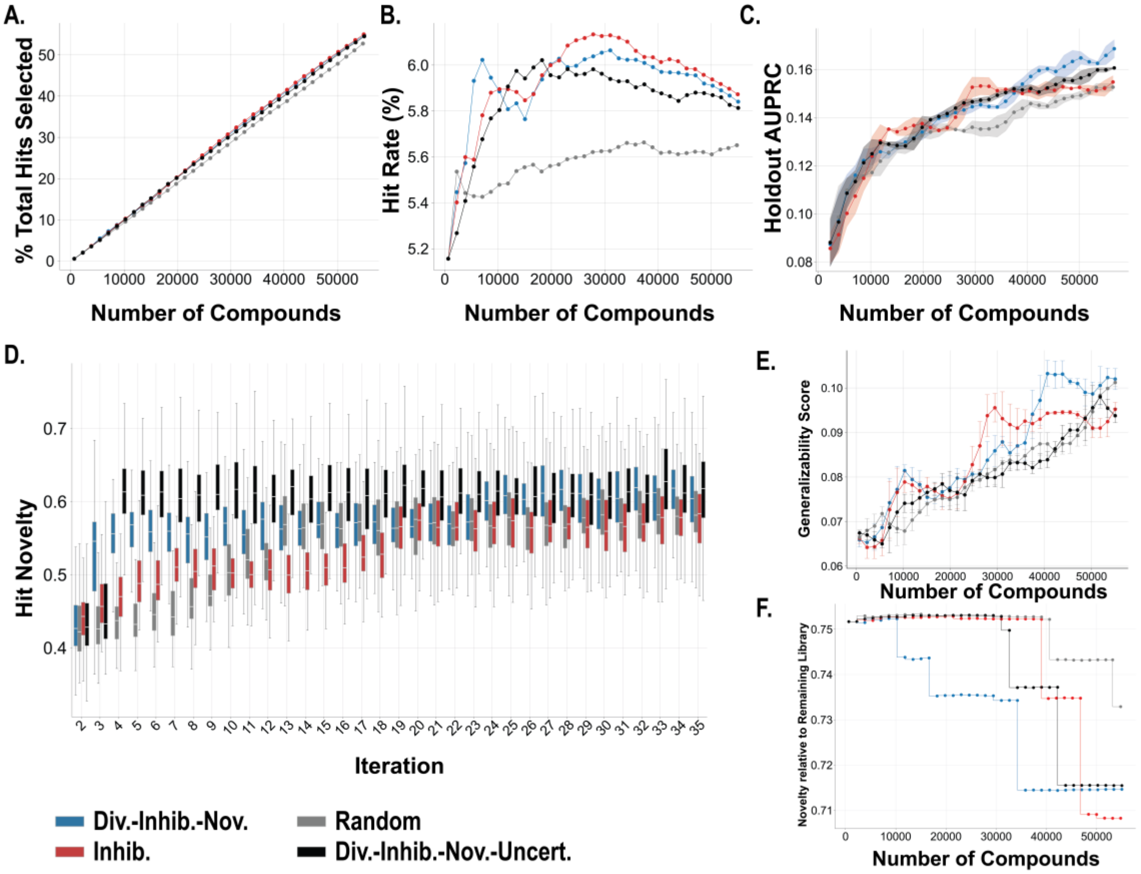


**Supplementary Figure S1. Simulation on *Mtb* HTS dataset with uncertainty. (A)** Percent total hits screened vs number of compounds **(B)** Number of active compounds screened per number of molecules screened ("hit rate"). **(C)** AUPRC vs number compounds. Each datapoint represents a new training batch selected based on the Random (gray), Diversity-Inhibition-Novelty (blue), Inhibition (red), or Diversity-Inhibition-Novelty-Uncertainty (black) acquisition functions. **(D)** Novelty (one minus the max. cosine similarity between MiniMol fingerprint of new hit and each hit from previous iteration) of active compounds screened at each iteration relative to the previous iteration. **(E)** Generalizability score (Methods) vs. number compounds for each condition. **(F)** Proxy metric for Remaining Library Novelty (average minimum cosine similarity of train molecules relative to remaining selection pool)

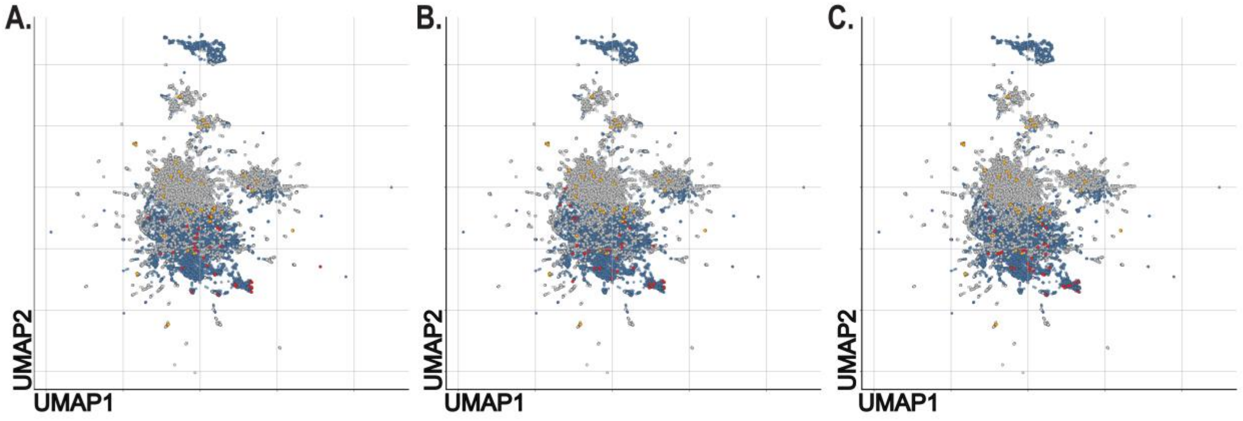


**Supplementary Figure S2. Subsampled AL hits for hit-controlled analysis.** UMAP of MiniMol representations of AL-selected non-hits (blue), investigator-selected non-hits (grey), subsampled AL-selected hits (red), and investigator-selected hits (orange) for AL hit control replicate 1 **(A)**, 2 **(B)** and 3 **(C)**.

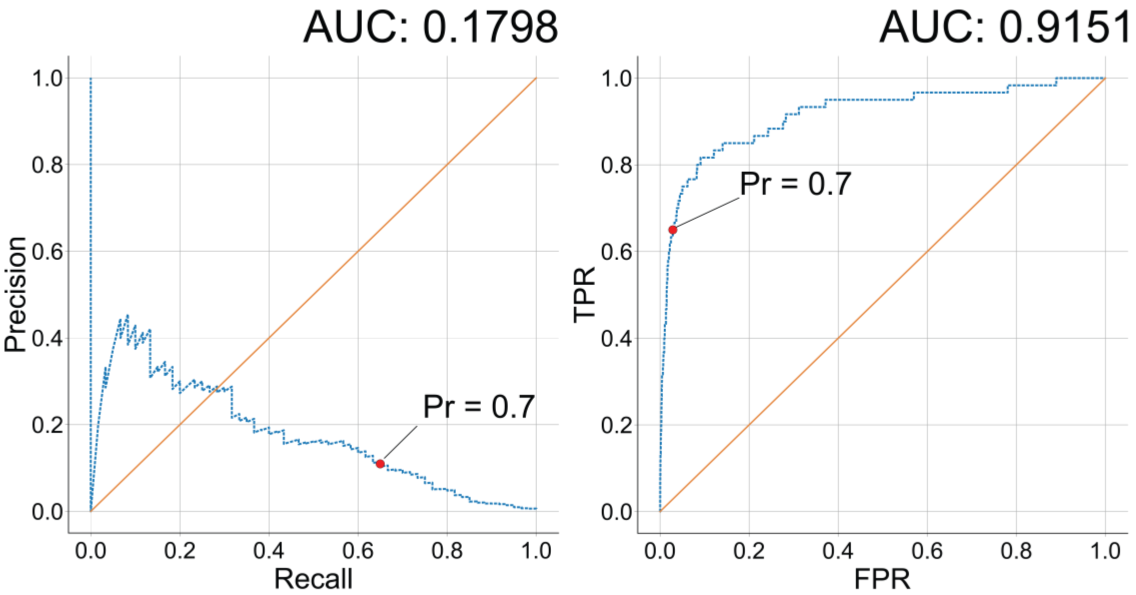


**Supplementary Figure S3.*B. burgdorferi* model used for virtual screening performance on holdout set.** Holdout set n = 10,540, number of hits = 60.

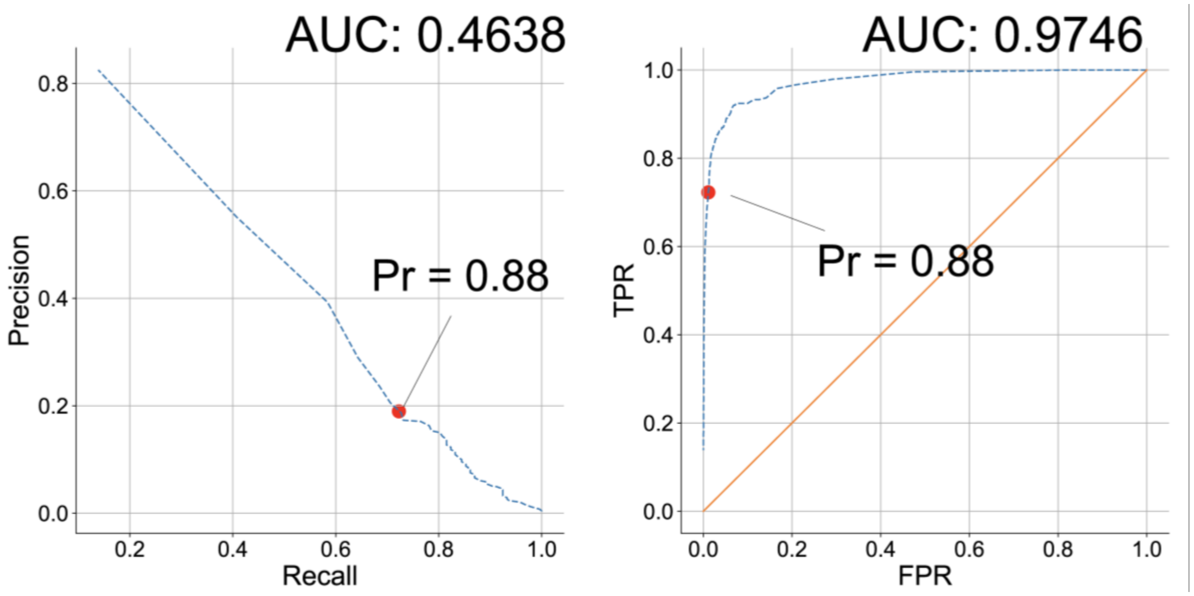


**Supplementary Figure S4**. ***S. aureus* model performance on holdout set.** 80/20 split train-test split. The final model incorporated the entire dataset (n = 327,979).

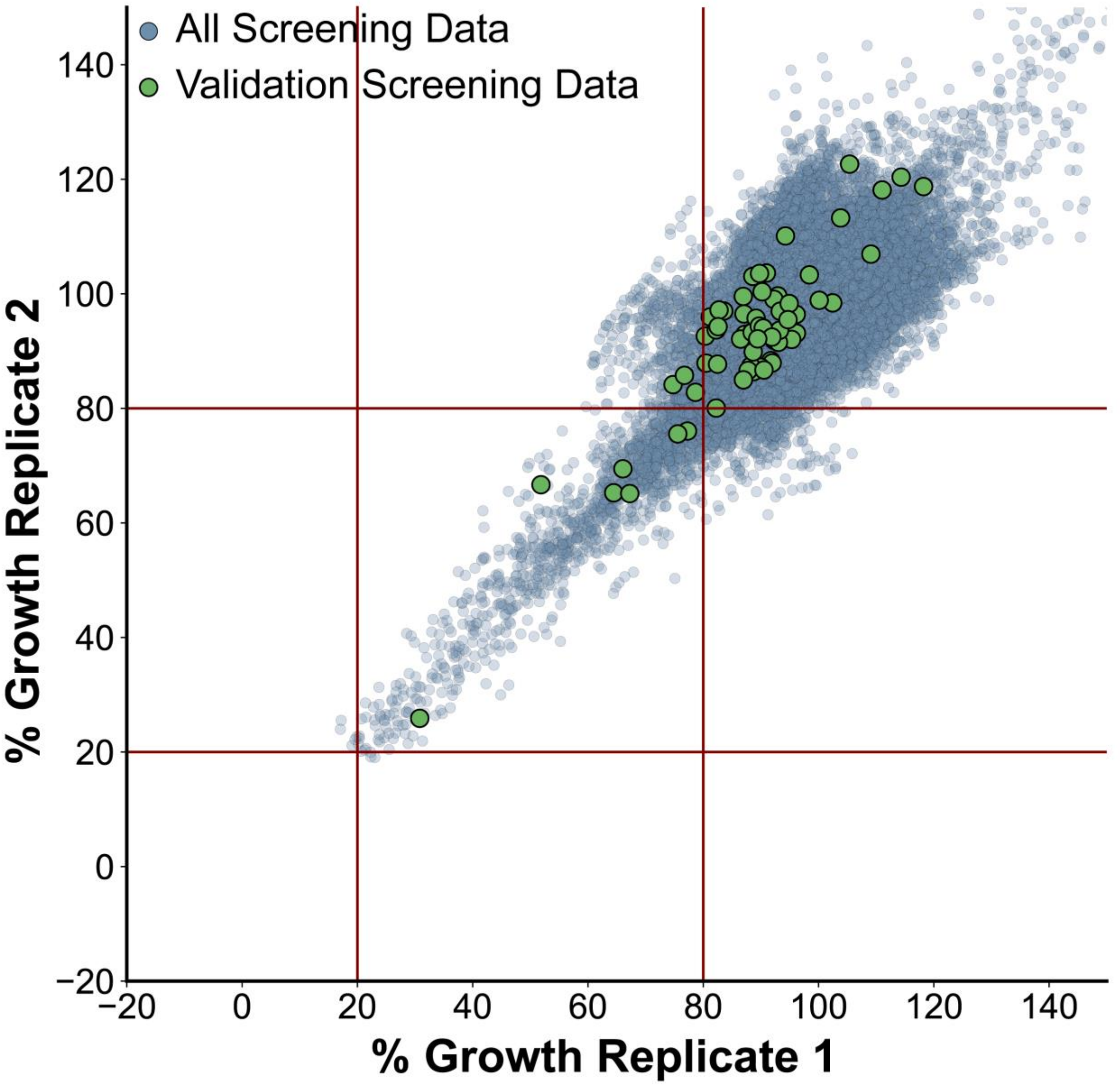


**Supplementary Figure S5.** Percent RFU relative to control replicates one and two plotted for all *B. burgdorferi* screening compounds (blue) and those for our validation screen (green) with growth inhibition of at least 80%.

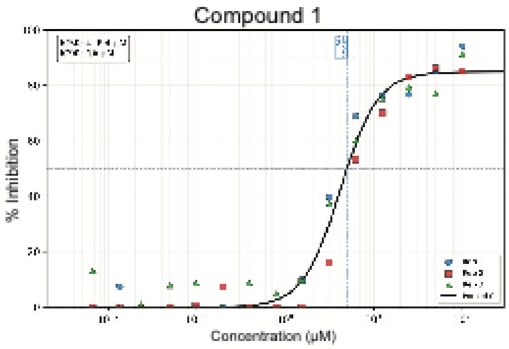
Compound 1
% Inhibition
Concentration (μM)

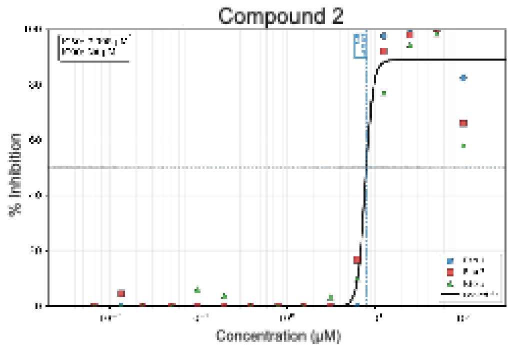
Compound 2
% Inhibition
Concentration (μM)

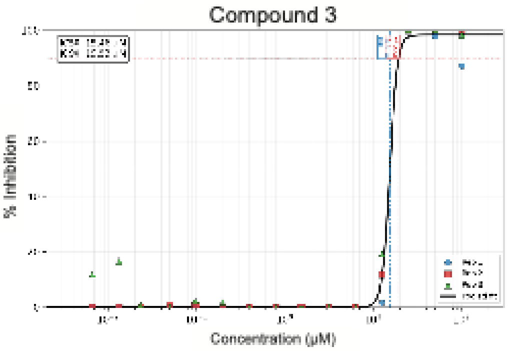
Compound 3
% Inhibition
Concentration (μM)

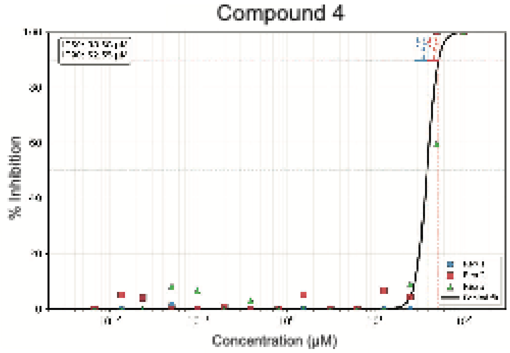
Compound 4
% Inhibition
Concentration (μM)

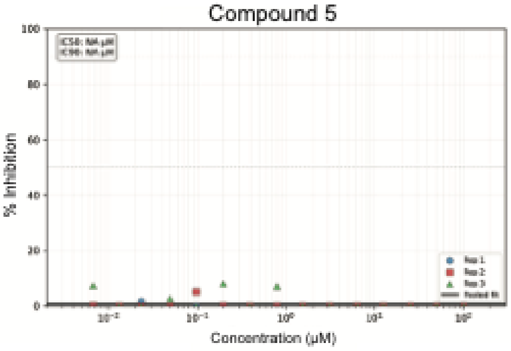
Compound 5
% Inhibition
Concentration (μM)

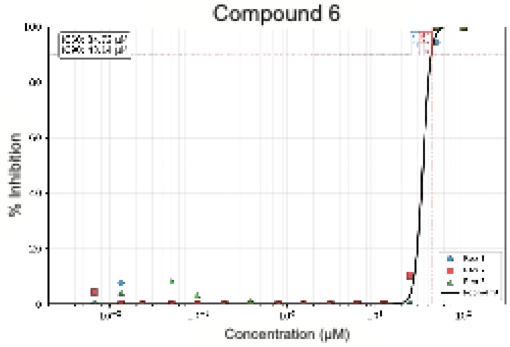
Compound 6
% Inhibition
Concentration (μM)

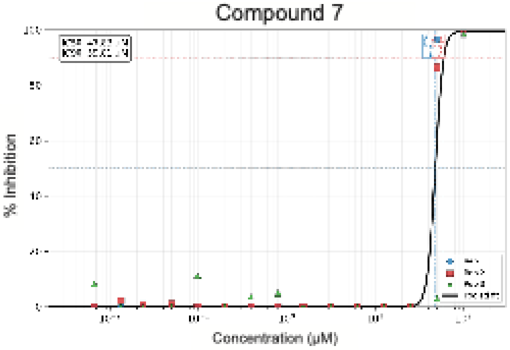
Compound 7
% Inhibition
Concentration (μM)

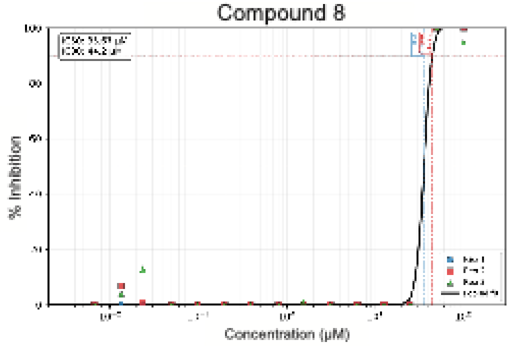
Compound 8
% Inhibition
Concentration (μM)

**Supplementary Figure S6. Dose response curves of 8 compounds identified by validation screen.** Compounds were tested in biological triplicate for inhibition in *B. burgdorferi*. $IC_{50}$ and $IC_{90}$ were determined by a fitted Hill curve.

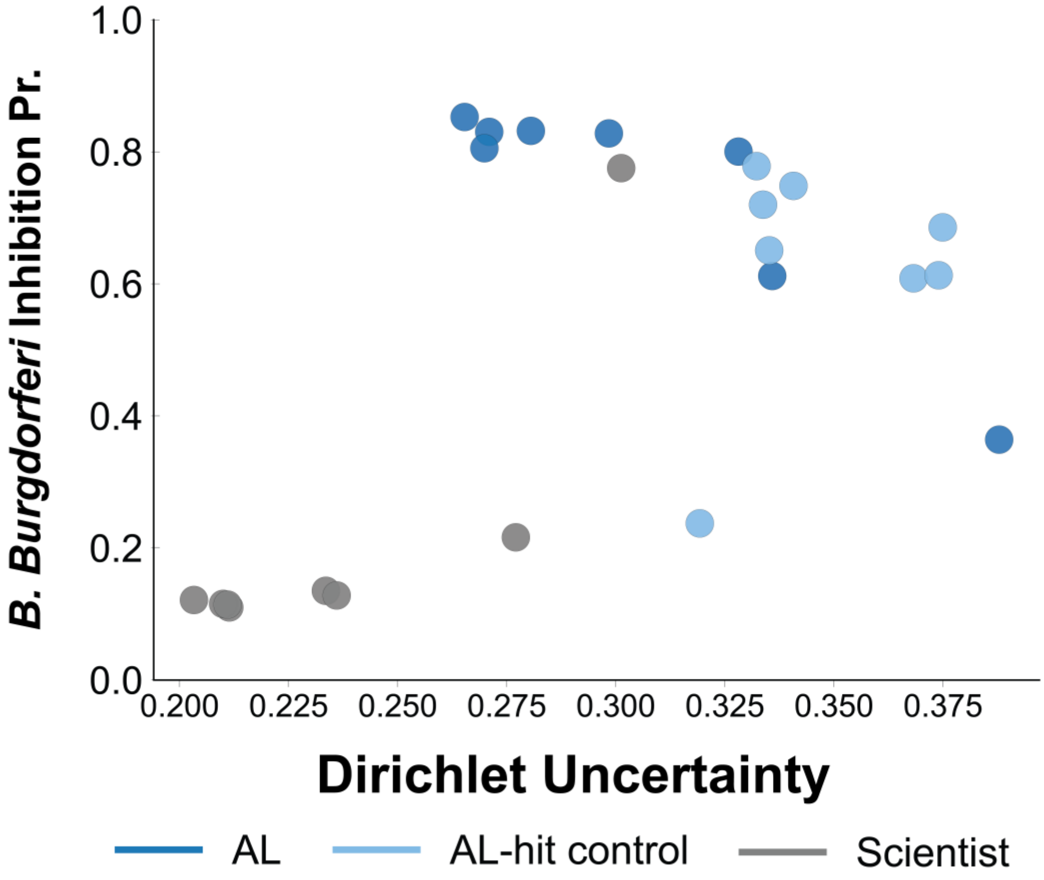


**Supplementary Figure S7.** *B. burgdorferi* inhibition probability vs. Dirichlet uncertainty for the AL-trained (dark blue), AL-trained controlled for hits (light blue), and investigator-curated-trained (grey)

| Fold | AUROC |
|---|---|
| 1 | 0.993 |
| 2 | 0.967 |
| 3 | 0.984 |
| 4 | 0.980 |
| 5 | 0.989 |
| mean | 0.983 ± 0.009 |

**Supplementary Table 1. *E. coli* model performance on holdout set.** Hit rate within the holdout set used for the *B. burgdorferi* model was insufficient for *E. coli* performance metrics (n = 2). *E. coli* model performance during K-fold cross validation during training is described.
